# Multiple modes of convergent adaptation in the spread of glyphosate-resistant *Amaranthus tuberculatus*

**DOI:** 10.1101/498519

**Authors:** Julia M. Kreiner, Darci Ann Giacomini, Felix Bemm, Bridgit Waithaka, Julian Regalado, Christa Lanz, Julia Hildebrandt, Peter H. Sikkema, Patrick J. Tranel, Detlef Weigel, John R. Stinchcombe, Stephen I. Wright

## Abstract

The selection pressure exerted by herbicides has led to the repeated evolution of herbicide resistance in weeds. The evolution of herbicide resistance on contemporary timescales in turn provides an outstanding opportunity to investigate key questions about the genetics of adaptation, in particular, the relative importance of adaptation from new mutations, standing genetic variation, or geographic spread of adaptive alleles through gene flow. Glyphosate-resistant *Amaranthus tuberculatus* poses one of the most significant threats to crop yields in the midwestern United States (1), with both agricultural populations and herbicide resistance only recently emerging in Canada (2, 3). To understand the evolutionary mechanisms driving the spread of resistance, we sequenced and assembled the *A. tuberculatus* genome and investigated the origins and population genomics of 163 resequenced glyphosate-resistant and susceptible individuals from Canada and the USA. In Canada, we discovered multiple modes of convergent evolution: in one locality, resistance appears to have evolved through introductions of preadapted US genotypes, while in another, there is evidence for the independent evolution of resistance on genomic backgrounds that are historically non-agricultural. Moreover, resistance on these local, non-agricultural backgrounds appears to have occurred predominantly through the partial sweep of a single haplotype. In contrast, resistant haplotypes arising from the midwestern US show multiple amplification haplotypes segregating both between and within populations. Therefore, while the remarkable species-wide diversity of *A. tuberculatus* has facilitated geographic parallel adaptation of glyphosate resistance, more recently established agricultural populations are limited to adaptation in a more mutation-limited framework.

**Significance:** While evolution is often thought of as playing out over millions of years, adaptation to new enviroments can occur in real time, presenting key opportunities to understand evolutionary processes. An important example comes from agriculture, where many weeds have evolved herbicide resistance. We have studied glyphosate resistant *Amaranthus tuberculatus*, a significant threat to crop yields in the midwestern US and Canada. Genome analyses showed that rapid evolution can either occur by “borrowing” resistance alleles from other locations, or by *de novo* evolution of herbicide resistance in a genetic background that was not previously associated with agriculture. Differences in recent evolutionary histories have thus favored either adaptation from pre-existing variation or new mutation in different parts of the *A. tuberculatus* range.

---

Glyphosate-resistant *A. tuberculatus* was first reported in Missouri in 2005, but has since been reported in 19 US states (1), with resistant biotypes harming corn and soybean yields (3, 4). Resistance to glyphosate in weed populations is widespread, likely as a result of the rapid adoption of and reliance on glyphosate weed control technology; 84% of soybeans, 60% of cotton, and 20% of corn grown in the US by 2004 carried transgenes for glyphosate resistance, despite Roundup Ready technology – the combination of glyphosate weed control with transgenic glyphosate resistance – only having been introduced eight years earlier (5). Agriculturally-associated *A. tuberculatus* weed populations emerged in Canada in the province of Ontario only in the early 2000’s, with glyphosate resistance following a decade later (2, 3). As with other herbicides, resistance in weed populations can evolve via substitutions at the direct target of glyphosate, 5-enolpyruvylshikimate-3-phosphate synthase (EPSPS), or by polygenic adaptation involving different loci in the genome (6–10). More often, glyphosate resistance in the genus *Amaranthus* can have an unusual genetic basis: amplification of the *EPSPS* locus (11–15). Gene amplification apparently evolved independently in two *Amaranthus* species (14, 16, 17), raising the possibility that it could have evolved multiple times independently within a single species, or even population (18). While glyphosate resistance has been studied from multiple angles (15, 19–23), the recent discovery of glyphosate-resistant A. *tuberculatus* in southwestern Ontario affords the unique opportunity to evaluate evolutionary origins of herbicide resistance, whether it has arisen through *de novo* mutation or standing genetic variation, and the role of gene flow in the recent spread of herbicide resistance in an agronomically important weed.

Native to North America, the dioecious, wind-pollinated *A. tuberculatus* has a history marked by the interaction of two lineages or subspecies (sensu Costea & Tardiff 2003 (19) and Pratt and Clark 2001 (20)), thought to have been diverging on either side of the Mississippi river until they were brought back into contact through human-mediated disturbance (21, 22). Morphological, herbarium, and microsatellite evidence point to an expansion of the western var. *rudis* subspecies range limits over the last 50 years, while the range of more eastern var. *tuberculatus* subspecies is thought to be stagnant and constrained to riparian habitats (22, 23). With the timing of the var. *rudis* expansion coinciding with the invasion of *A. tuberculatus* into agricultural environments, var. *rudis* is hypothesized to be a predominant driver of this agricultural invasion (23).

We assembled a high-quality reference genome for *A. tuberculatus* from a single individual from 58 Gb (approx. 87X genome coverage) long-read data collected on the Pacific Biosciences Sequel platform using 15 SMRT cells. After assembly, polishing, and haplotype merging, the reference genome consisted of 2,514 contigs with a total size of 663 Mb and an N50 of 1.7 Mb (see Sup Table 1 for details). Our final genome size is consistent with recent cytometric estimates of 676 Mb (SE=27 Mb) for *A. tuberculatus (24)*. The new reference included 88% of the near-universal single copy orthologs present in BUSCO’s Embryophyta benchmarking dataset, with 6% marked as duplicates (25). For chromosome-scale sweep scan analyses, we further scaffolded our contigs onto the fully resolved *A. hypochondriacus* genome (26), resulting in 16 final pseudomolecules (which included 99.8% of our original assembly, see methods) for population genetic analyses.

We resequenced whole genomes of 163 individuals to about 10X coverage from 19 agricultural fields in Missouri, Illinois, and two regions where glyphosate resistance has recently appeared in Ontario: Essex County, an agriculturally important region in southwestern Ontario, and Walpole Island, an expansive wetland and First Nation reserve with growing agricultural activity (Fig. 1B). Populations from the Midwest (Missouri and Illinois) had been previously assayed for glyphosate resistant phenotypes, and from qPCR of genomic DNA it was found that resistance was predominantly conferred through *EPSPS* copy number amplification (11). Populations from Walpole Island and Essex County in Ontario were sampled in 2016 after reports from farmers as going uncontrolled. We also sampled 10 individuals from riparian habitats in Ontario near Walpole Island & Essex county, as a non-agricultural, natural Canadian comparison (Fig. 1B). Genomewide diversity in *A. tuberculatus* is quite high, even relative to other wind-pollinated outcrossers (27), with neutral diversity (mean pairwise difference) at four-fold degenerate sites being 0.041. Glyphosate resistance in the sampled agricultural fields ranged from 13% to 88%, based on greenhouse trials (Fig. 1B; see Methods). Plants from natural populations in Ontario had no glyphosate resistance.

**Fig. 1.**
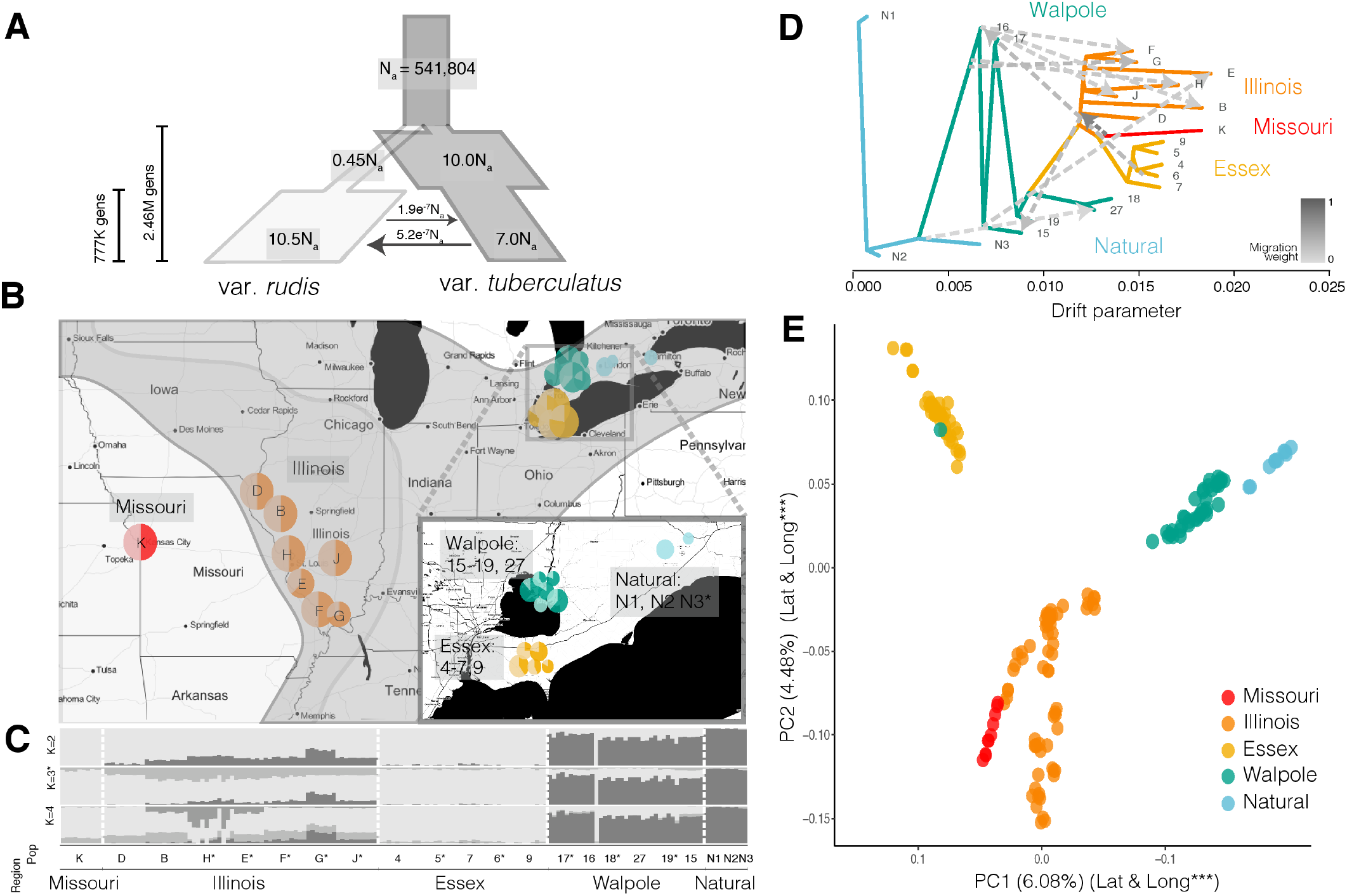
Population structure and demographic history in *Amaranthus tuberculatus*. (*A*) Demographic model of the two *A. tuberculatus* subspecies (N_a_ = ancestral effective population size, with migration and other effective population size estimates scaled accordingly). (*B*) Geographic map of phenotyped and sequenced populations with glyphosate resistance in US and Canada. Natural populations are non-agricultural, Canadian populations without glyphosate resistance. Population are color coded by region (maintained throughout text). Dark portions of each pie chart indicate proportion of resistant individuals. Inset shows closeup of agricultural and natural Ontario populations. Historical range limits of the species are indicated on the map, with the same coloration as for the demographic model – *A. tuberculatus* var. *rudis* in light grey to the west, and *A. tuberculatus* var. *tuberculatus* in dark grey to the east (adapted from Sauer 1955). (*C*) STRUCTURE plot of admixture across regions and populations from west (left) to east (right), with most likely number of clusters (K) = 3. The darkest ancestry group corresponds to var. *tuberculatus*, and the lightest corresponds to var. *rudis*. (*D*) *treemix* results showing the maximum likelihood of relatedness between populations based on allele frequencies, with population labels at the tip of each branch, dashed arrows indicating the amount of migration between populations, and the drift parameter reflecting the amount of genetic drift that has occurred between populations. (*E*) PCA of all individuals, with both PC1 and PC2 significantly relating to both longitude and latitude (See also Table S2).

## Demography of *Amaranthus tuberculatus*

To dissect the demographic context of convergent adaptation to glyphosate, we characterized genome-wide patterns of population structure, demography, and differentiation. Population structure, demographic modelling (Fig. 1), and phenotypic characterization confirmed the presence of the two previously hypothesized ancestral lineages, *A. tuberculatus* var. *rudis* and *A. tuberculatus* var. *tuberculatus (22, 23)*. Population structure and investigations of the genome-wide proportion of introgression (*f* statistic (28)) largely reflected previous accounts of the historical range limits (22): natural Ontario populations had the diagnostic indehiscent seed phenotype and were genetically homogeneous for ancestry of the var. *tuberculatus* lineage, Missouri samples were homogeneous for the var. *rudis* lineage, while Illinois, a region of sympatry in the historical range of the two subspecies, showed signs of introgression from var. *tuberculatus* (*f* [95% CI]= 0.1342 [0.126, 0.143], using Missouri as a reference) *(22)* (Fig. 1B,C). Genetic differentiation (F_ST_) between individuals with ancestry homogenous for different lineages at K=2 was 0.212, on par with or greater than that between congeners (29). Moreover, both longitude and latitude significantly explained both PC1 and PC2 of the SNP matrix (Sup Table 2), with PC1 separating var. *rudis* and var. *tuberculatus* ancestry, and PC2 separating Canadian and American accessions (Fig. 1E). These patterns resulted in a PCA representation that, with few exceptions, reflected the geography of our samples. The most likely *tuberculatus-rudis* demographic model was one of secondary contact, with var. *rudis* having undergone a bottleneck followed by a dramatic expansion, which may be indicative of this subspecies’ rapid colonization of agricultural fields across North America (Fig. 1A).

## Demographic origins of Canadian agricultural populations

Analyses of agricultural populations in Ontario, which have only recently become problematic, shed new light on the demographic source of the *A. tuberculatus* agricultural invasion. Populations from Essex county fell completely within the var. *rudis* cluster, with a *treemix* model indicating that Essex populations were derived from the most western, Missouri population (Fig. 1E), the source of almost the entire Essex genome (*f* =0.996, 95% CI 0.985–1.0). Furthermore, while Essex grouped with Walpole and Natural populations on PC2, it was found at the other end on PC1, more different from Canadian populations than even the most geographically distant Missouri population (Fig. 1B). These patterns of population structure were distinct from the continuous gradient of west-east ancestry previously reported (23), and supports the hypothesis that glyphosate-resistant *A. tuberculatus* was introduced to Ontario through seed-contaminated agricultural machinery (2, 3) or animal-mediated seed dispersal (30).

In contrast to Essex as a likely introduction of a preadapted genotype to a new locale, populations from Walpole Island, where glyphosate resistance was first reported in Ontario (2), were mainly of the native, eastern var. *tuberculatus* type (Fig. 1). However, the convergent evolution of var. *tuberculatus* into agricultural fields may not be solely the result of *de novo* mutations. Populations from Walpole Island showed signs of introgression from var. *rudis* (*f* =0.225, 95% CI 0.215–0.236), while *treemix* indicated that Walpole may be a hotspot for gene flow, with 9/10 total migration events across the tree involving Walpole (explaining an additional 2.5% of SNP variation compared to a migration-free model) (Fig. 1C). Thus, both adaptive introgression from the western var. *rudis* clade and/or *de novo* adaptation from local natural populations could be playing a role in the evolution of resistance and adaptation to agricultural environments in Walpole.

Despite the considerable level of var. *rudis* introgression into Walpole, these populations were similarly differentiated from nearby natural populations homogenous for var. *tuberculatus* ancestry as they were from comparably admixed populations in Illinois (F_ST (Walpole-Nat)_=0.0286; F_ST (Walpole-Illinois)_=0.0284; Fig. S1). This, along with the tight clustering of Walpole and Natural populations in the PCA and structure analyses, implies that Walpole populations experienced strong and rapid local adaptation to agricultural environments upon its conversion from wetland, which may have been facilitated by introgression from var. *rudis*. We therefore sought to find genes that were highly differentiated between Walpole and Natural populations, putatively involved in agricultural adaptation. A GO enrichment test for the top 1% of genes with excess differentiation between Walpole and natural populations identified genes with monooxygenase and oxidoreductase molecular function, with histone methylation biological function, and several protein classes involved in transport and amino acid/protein modification (Fig. 2A). Of particular interest were two enriched GO categories important in metabolic, non-target site resistance: oxidoreductases, which include cytochrome P450s, and peptidyl-amino acid modifiers, which include glycosyltransferases (7). CYP450s and glycosyltransferases work consecutively to detoxify herbicides in plant cells by catalyzing hydroxylation and glc-conjugation (31). Beyond being highly differentiated between Walpole and Natural populations, an analysis of the maximum copy number in 1 kb windows within each CYP450 and glycosyltransferase gene (we used large windows to control for heterogeneous paralog/ortholog degeneration) revealed that both gene families have expanded copy number in Walpole agricultural populations compared to natural populations (CYP450s; *F*_1,3239_=6.65, *p*=0.0099; glycosyltransferases; *F*_1,3004_=3.313, *p*=0.0688) (Fig. 2B). Despite this widespread pattern across 69 CYP450 and 64 glycosyltransferase genes, only 2 CYP450s and 3 glycosyltransferases were significantly positively correlated with our phenotypic rating of glyphosate resistance, after Holm’s correction for multiple testing (Sup Table 3). A possible explanation is that the copy number expansion of these gene families confers resistance to herbicides other than glyphosate, and is a result of the transition from natural to agricultural habitats.

**Fig. 2.**
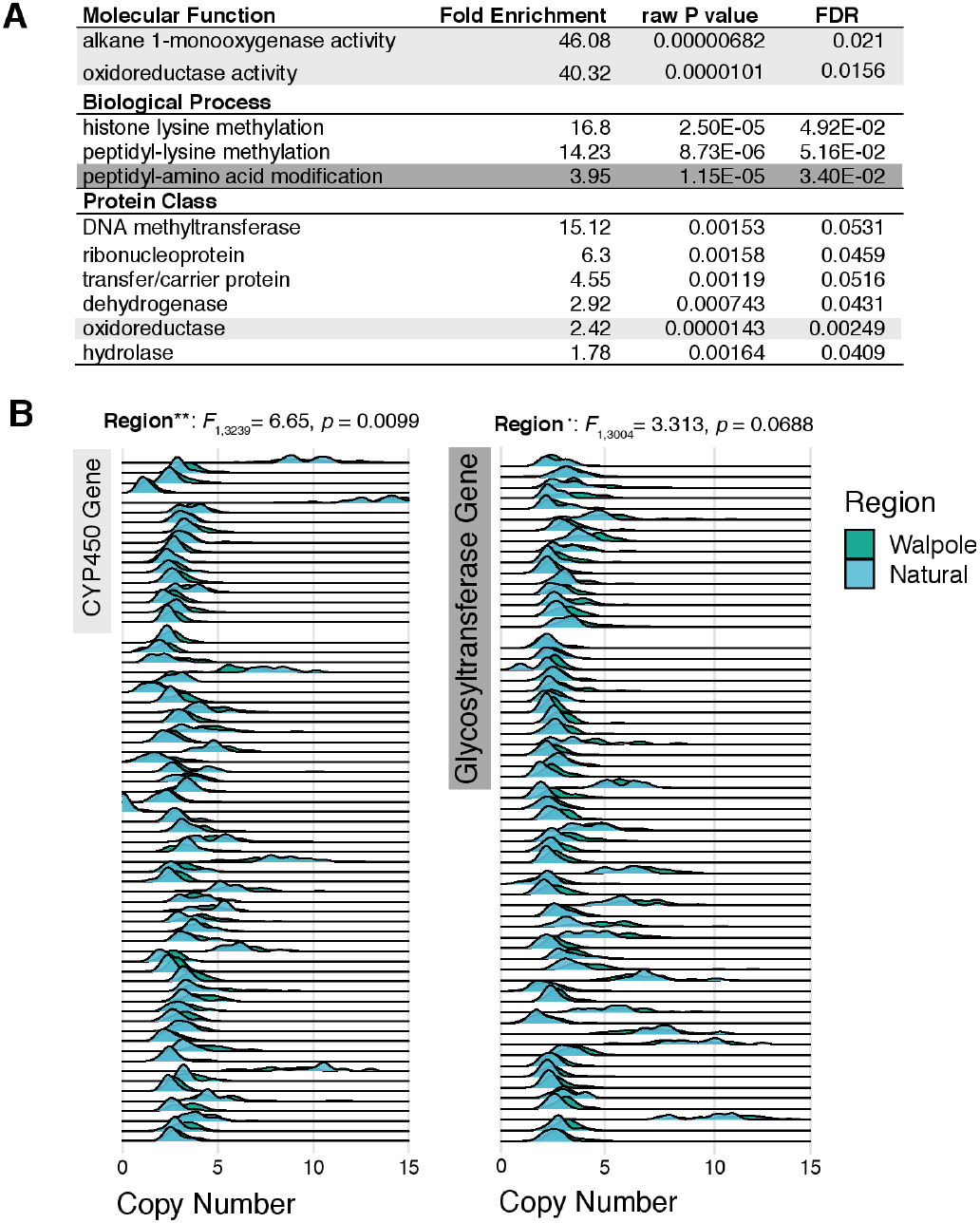
Enrichment and expansion of CYP450 and glycosyltransferase gene families in the transition from natural to agricultural in Walpole. (*A*) GO categories that were significantly enriched in an analysis of the 99^th^ percentile of Fst outliers between Walpole and Natural populations. Light grey indicates GO categories that include CYP450s, dark grey indicates the category that includes glycosyltransferases. (*B*) Evidence for copy number expansion of both CYP450 and glycosyltransferase genes in Walpole relative to Natural populations. Genes that significantly relate to phenotypic resistance are shown in **Fig. S3**.

## Genetic mechanisms of glyphosate resistance

Two major evolutionary paths to glyphosate resistance are amplification of wild-type *EPSPS* and non-synonymous mutations in *EPSPS* that make the enzyme resistant to glyphosate inhibition. To better understand the genetic mechanisms underpinning glyphosate resistance, we investigated how variation in resistance relates to these two classes of *EPSPS* mutations. Using our genomic data to quantify copy number (see methods), we found that of 84 individuals assayed in the greenhouse as resistant (resistance => 2/5 rating; Fig. 3), 60 (71%) had elevated *EPSPS* copy number (> 1.5; as in (13)). Almost 26%(22/83) of individuals assayed as susceptible had an *EPSPS* copy number > 1.5 (compared to 15% or 13/88 individuals for a >2 cutoff). Apart from errors in phenotyping or copy number estimation, this implies that at least intermediate copy number amplification alone may not always be sufficient for resistance, e.g., if amplified copies are not properly expressed. While *EPSPS* amplification was most frequent in the Midwest (83% [33/40] of resistant individuals, compared to 70% [16/23] in Walpole and 52% [11/21] in Essex), copy number in resistant individuals was on average almost twice as high in Walpole (⊠ 9 copies on average, compared to 5 in the Midwest, and 4 in Essex). Previous estimates of *EPSPS* copy number in resistant *A. tuberculatus* were up to 17.5 copies relative to diploid susceptibles (11); we found two individuals in Walpole with an estimated 29 copies (Fig. 3). A regression of resistance onto copy number was significant in all three geographic regions (Walpole p=2.6e-07; Essex p=0.002; Midwest p=3.5e-06), explaining 48% of the variation in resistance in Walpole, but only 23% and 27% in Essex and the Midwest. In these latter two regions, however, an additional 10% of variation was explained by a non-synonymous substitution at codon 106, the most common and well-characterized genetic mechanism of glyphosate resistance across species outside of the genus *Amaranthus* (1), in this instance causing a change from proline to serine (Fig. 3).

**Fig. 3.**
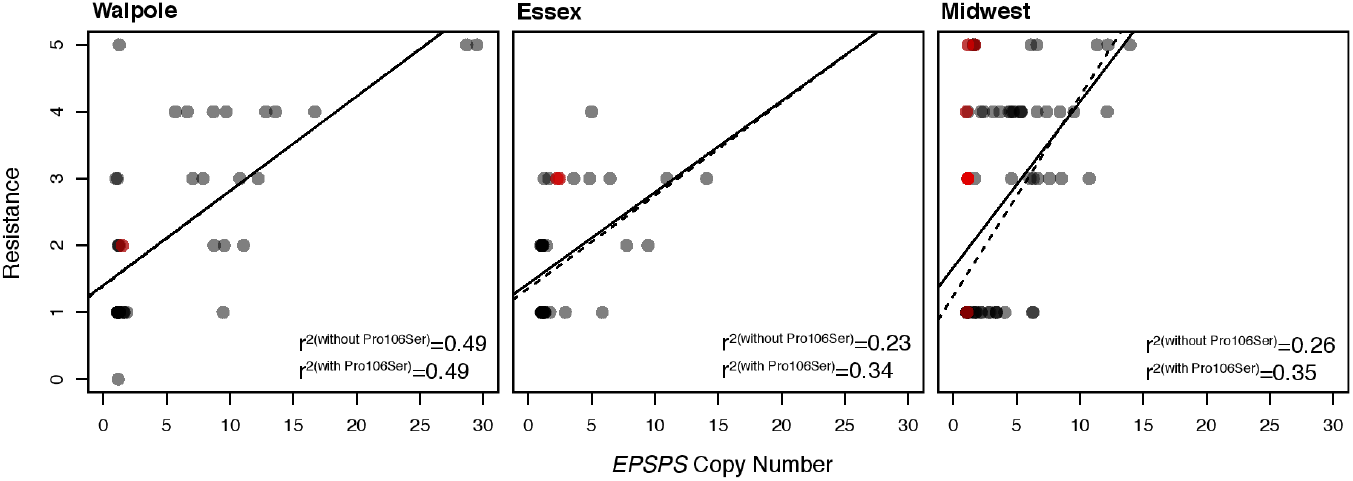
*EPSPS* copy number variation among individuals, and its relationship with resistance. *EPSPS* copy number significantly explains phenotypic resistance within each agricultural region (solid linear regression line) (Walpole p= 2.6e-07; Essex p=0.002; Midwest p=3.5e-06), with P106S substitution in *EPSPS* (red individuals) increasing explained resistance in Essex and the Midwest, but not in Walpole (dashed linear regression line).

The presence of two types of target-site resistance mechanisms, copy number increase and non-synonymous mutation at a critical codon, implies parallel evolution of the resistance phenotype through independent genetic pathways. While the well-known P106S non-synonymous mutation can account for some of the resistance unexplained by copy number increase alone, other uncharacterized non-target site mechanisms are likely contributing as well and thus providing a further path to convergent evolutionary outcomes. In addition to shedding light on the prevalence of different resistance mechanisms, the population genomic data allowed us to determine whether our most prevelant genetic mechanism, namely the *EPSPS* gene amplification, arose multiple times.

## Genetic origins of the *EPSPS* amplification

Our chromosome-scale genome assembly provided a unique opportunity to determine the structure and genomic footprint of selection around the amplified *EPSPS* locus in different populations. Across all populations, copy number increase was not restricted to the 10 kb *EPSPS* gene—individuals identified to have increased copy number at *EPSPS* also had a correlated increase in the mean and variance of copy number for up to 6.5Mb (23.5 – 30 Mb on chromosome 5) of the reference genome, encompassing 108 genes (Fig. S2).

Characterizing signals of selection for a high copy number region can be challenging. First, typical population genetic statistics ignore potential variation among gene copies that are collapsed into a single haplotype. Ideally, phasing of a multicopy region would allow for full resolution of the SNP differences within and between haplotypes. However, very recent gene amplification is expected to limit SNP variation among amplified copies, and will also hinder phasing approaches from short read data. Second, variation may not be recognized because of allelic dropout of low-copy variants. Analysis of the relationship between EPSPS copy number and homozygosity in our dataset suggested that higher-copy haplotypes did not feature more SNP variation than lower-copy haplotypes, suggesting that generally few new mutations distinguish among amplified copies (Fig. S3). To control for the possibility of residual SNP differences that exist among gene copies and/or for allelic dropout, we created a consensus haploid sequence by random downsampling to one allele per heterozygous site.

While the *EPSPS*-related amplification showed the strongest selective signal on all of chromosome 5, we found distinct selective sweep patterns in the different agricultural regions. We ran *Sweepfinder2 (32, 33)* across chromosome 5 to identify focal windows with a site-frequency spectrum particularly skewed by selection (while controlling for recombination) relative to genomewide 4-fold degenerate sites. *Sweepfinder2* estimated the strongest amplification-related sweep signal in Walpole. In contrast to Essex or the Midwest, the top 1% of apparently selected 10 kb windows on chromosome 5 were localized to the amplified *EPSPS* region in Walpole (Fig. S4). Moreover, there was a marked reduction in genetic diversity (mean pairwise differences) around *EPSPS*, as well as elevated differentiation (F_st_) and extended haplotype homozygosity (XP-EHH score (34)) in Walpole individuals with *EPSPS* amplification, implying a hard selective sweep, but not in Essex or Midwest individuals with increased *EPSPS* copy number (Fig. 4).

**Fig. 4.**
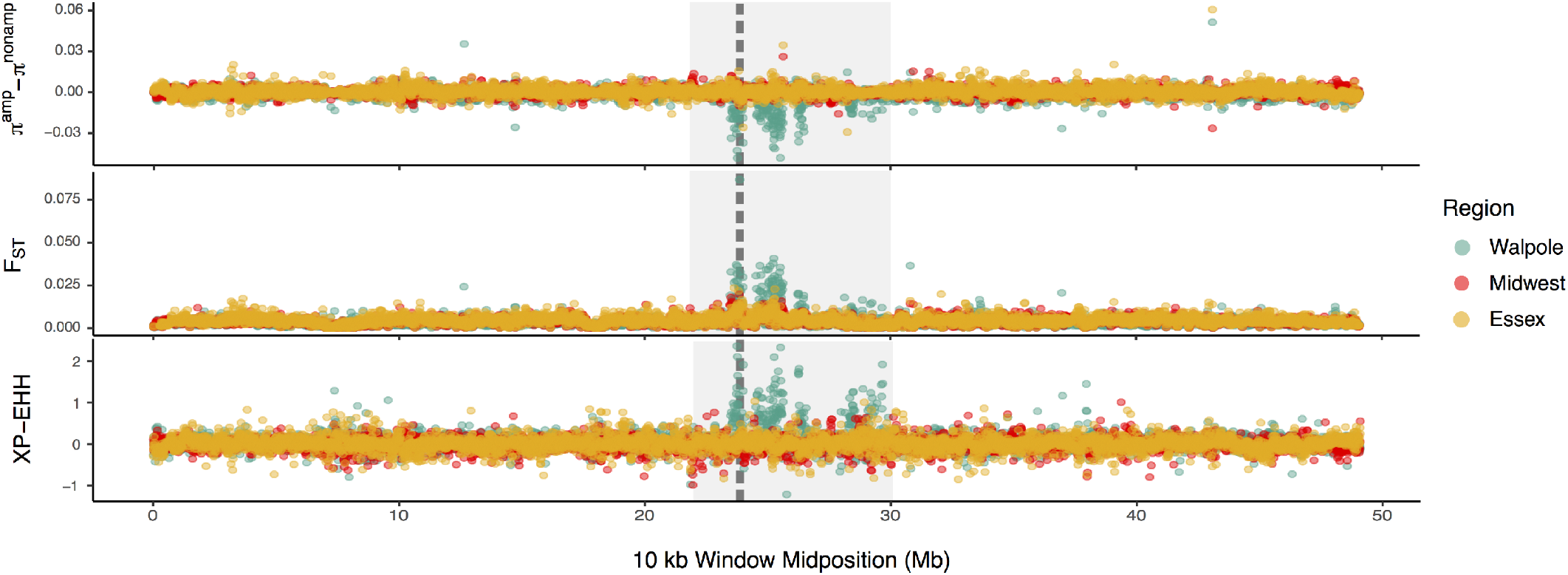
Population genetic signals of selection related to copy number increase around *EPSPS* on chromosome 5. The deficit of diversity (*top*), relative differentiation (*middle*), and difference in extended haplotype homozygosity (XP-EHH (34)) (*bottom*) is compared between amplified and non-amplified individuals in each agricultural region. *EPSPS* is delimited by the vertical grey dashed line, while the *EPSPS*-linked region undergoing amplification is shown by the light grey box, spanning 23.5 to 30 Mb on chromosome 5.

These differences in the extent of the amplification-related sweep signals across agricultural regions may be a consequence of how often *EPSPS* amplification has evolved; a hard sweep would be indicative of it having arisen only once, while soft sweeps would point to multiple origins (35–38). To investigate this further, we mapped *EPSPS* copy number onto a maximumlikelihood haplotype tree produced from SNP variants in *EPSPS*, and compared the phylogeny with phenotypic resistance and non-synonymous target-site resistance status (Fig. 5). Indeed, the agricultural regions differed in the inferred number of independent copy number increases. Whereas there appears to have been only one amplification event in Walpole, Essex haplotypes of individuals with copy number increases are interspersed with susceptible haplotypes, both within and between populations. Similarly, haplotypes from Midwest individuals with *EPSPS* amplification are distributed across the gene tree, although some local populations show clustering indicative of a local hard sweep, implying independent evolutionary origins among populations and occasionally within populations in the Midwest (Fig. 5).

**Fig. 5.**
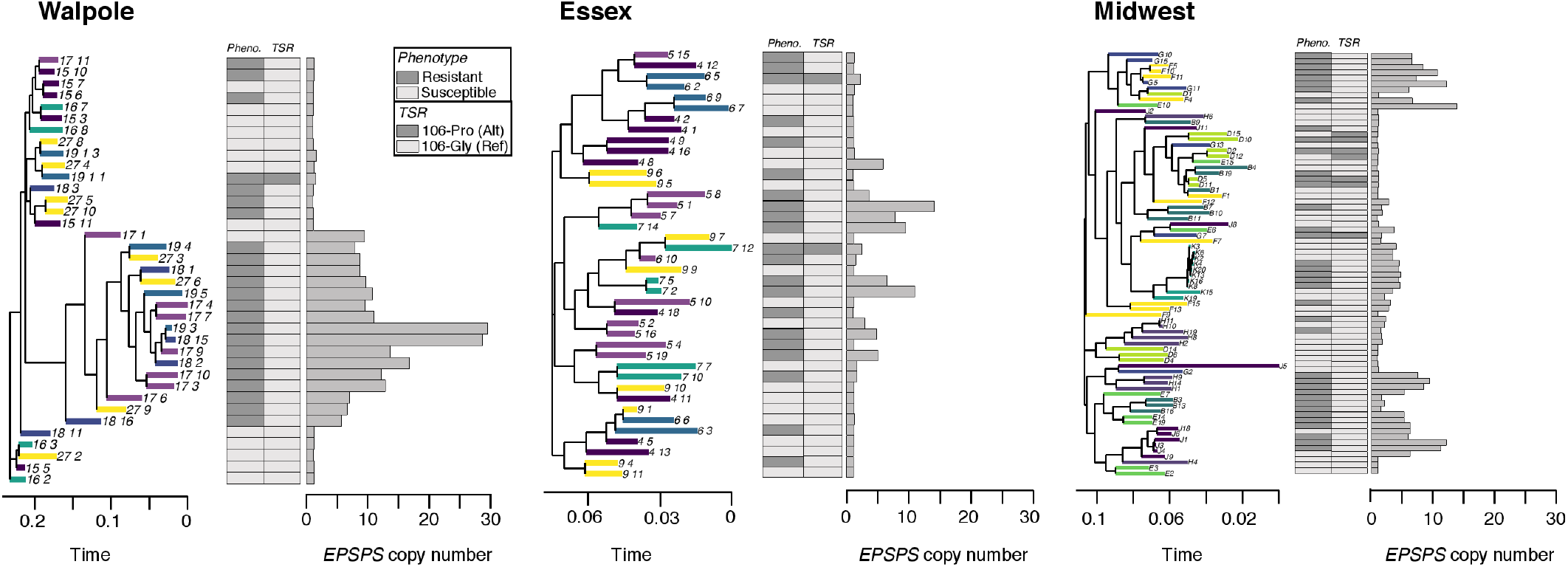
Diversity of *EPSPS* amplification origins across agricultural regions. For each agricultural region, we show a haplotype treeof the 1Mb region surrounding *EPSPS* (with tips coloured by population of origin), alongside a barplot of *EPSPS* copy number, and a matrix of phenotypic resistance and targetsite-resistance status for the Gly-106-Pro mutation.

Together, these analyses suggest that gene amplification has occurred multiple times independently to different extents across the geographic range. However, it is possible that recombination and *de novo* mutation after amplification have contributed to the apparent soft sweep signal. To further test for such a scenario, we looked at the similarity in the copy number profiles of the *EPSPS* region, which should be independent of any possible artifacts due to minority allele drop out in resequencing data. The copy number profiles of the amplified region varied considerably across our samples, and especially across agricultural regions (Fig. 6A), consistent with multiple independent amplification events. To quantify this, we calculated the correlation in normalized sequencing coverage in the 1 Mb chromosomal segment surrounding *EPSPS* between all possible pairs of individuals with copy number increases (Fig. 6B). In agreement with our polymorphism-based inferences, the two Canadian regions showed very different patterns; coverages in different Walpole individuals were very highly correlated (average of Spearman’s ρ = 0.95), suggesting the spread of a single amplification haplotype through a hard selective sweep. In contrast, there was much a lower average correlation across all Essex individuals regionwide (ρ= 0.56), and this was the case even when looking at the average within-population correlations rather than the single region-wide average (within-pop average e.g.: ρ = 0.54 and 0.61), suggesting different haplotypes had independently experienced copy number increases (Fig. 6). Similar to Essex, there appeared to be multiple amplification haplotypes in the Midwest (average for all individuals, ρ= 0.47), but within-population correlations were higher, consistent with hard (ρ= 0.94, 0.95, 0.93) or soft sweeps (ρ=0.66, 0.74, 0.75) (Fig. 6).

**Fig. 6.**
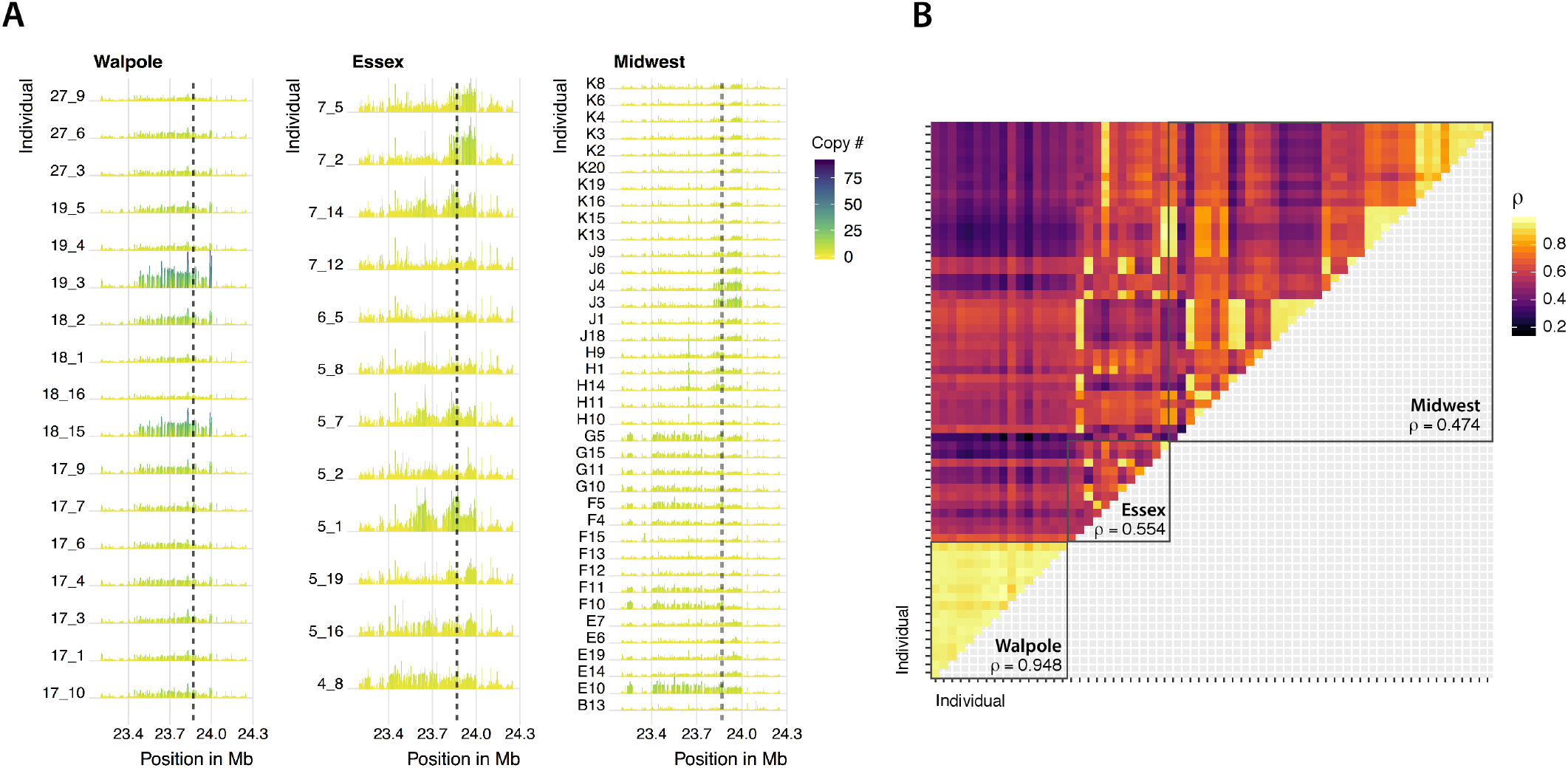
Similarly of the *EPSPS* amplification inferred from copy number variation around the *EPSPS* gene. (*A*) Normalized sequencing coverage for 1Mb around EPSPS (locus delimited by dashed line). Only individuals with at least 1.5X coverage relative to susceptible individuals were considered. (*B*) Pairwise Spearman’s correlation coefficients over the same 1-Mb *EPSPS* region for individuals shown in (A). Mean coefficient across all individuals within each agricultural region indicated. Order of individuals within each region as in (A), from bottom-top & right-left.

The patterns of genetic differentiation and similarity in amplification profiles among agricultural regions helped us to distinguish between modes of adaptation, the evolutionary mechanisms by which glyphosate resistance has spread, and the extent of constraint on this particular genetic pathway. Although the Walpole population showed signs of admixture from var. *rudis*, Walpole individuals were clearly differentiated at *EPSPS* from both Essex and Midwest individuals (Fig. S5). Moreover, copy number profiles were almost perfectly correlated within Walpole, but low with Essex and the Midwestern individuals (Fig. 6B). This suggests that glyphosate resistance in Walpole evolved independently, likely from selection on a *de novo* amplification event, although we do not know whether the amplification occurred in Walpole, or whether this allele was introgressed from an unsampled population. In Essex, the lack of within region correlation in *EPSPS* copy profiles and sporadic high correlations with individuals from different Midwestern populations (Fig. 6B), suggest multiple independent amplification events. Together, with the lack of genetic differentiation between Essex and Midwest (both genome wide and on all of chromosome 5, including EPSPS Fig. S5), this suggests that Essex was either directly colonized by a diverse glyphosate resistant population from the Midwest, or that a prior glyphosate susceptible population in Essex was replaced by glyphosate resistant individuals from the Midwest.

In summary, we have found multiple modes of convergent evolution underlying the spread of glyphosate resistance in North American *A. tuberculatus* populations. There is evidence for a single *EPSPS* amplification event that gave rise to the resistant populations in Walpole and that is different from amplification events in populations from another Canadian region, Essex, and from populations in the US Midwest, where glyphosate resistance is older than in Canada. In contrast to the hard sweep in Walpole, glyphosate selection has left only soft selective sweep signals in the Midwest, because different haplotypes were amplified independently. Together with our analyses of population structure and demographic history, these results suggest that evolution on the more agriculturally-naive *A. tuberculatus* var. *tuberculatus* background occurred in a mutation-limited framework, relying on evolutionary rescue via *de novo* mutation. In contrast, as suggested by Kreiner and colleagues (39), a longer history of temporally and geographically fluctuating selection for glyphosate resistance on the *A. tuberculatus* var. *rudis* background in the Midwest maintains multiple independent amplification haplotypes both within and among populations, some of which appear to have spread to Essex via gene flow. Therefore, genetic background and duration of selection interact to determine whether adaptation remains constrained toamutation-limited framework.

A practical outcome of this work is that it informs on the scale of management that is needed to control herbicide resistance. Specifically, we suggest that with glyphosate resistance spreading across the range through seed translocation and independent adaptation, management efforts should be broadened to encompass both regional seed containment and local integrative control of herbicide-resistant weeds. We are faced with an additional challenge—that historically non-weedy lineages can adapt to an agricultural environment on rapid, contemporary timescales—calling for more consideration of how to prevent seemingly benign weeds from becoming problematic.

## Methods

### Plant Collections

Seeds were collected from Midwestern populations in 2010 (11), and from Ontario natural populations and agricultural fields in the fall of 2016. Agricultural fields in which *A. tuberculatus* appeared to be poorly controlled were sampled, biasing the collection towards populations with high levels of glyphosate resistance. These do not necessarily represent levels of resistance in a random sample.

### High Molecular Weight DNA Extraction

High molecular weight (HMW) DNA was extracted from the leaf tissue of a single 28-day-old female *A. tuberculatus* plant from the Midwest using a modified version of the Doyle and Doyle nuclei isolation protocol (40). Nuclei isolation was carried out by incubating 30 g of ground leaf tissue in a buffer comprising tris(hydroxymethyl)aminomethane, potassium chloride, ethylenediaminetetraacetic acid, sucrose, spermidine and spermine tetrahydrochloride (Sigma-Aldrich, MO). The homogenate was subsequently filtered using miracloth and precipitated by centrifugation. G2 lysis buffer, RNase A, and Proteinase K (Qiagen, Venlo, Netherlands) were then added prior to an overnight incubation at 50°C, followed by centrifugation at 4°C. The supernatant containing the DNA solution was added to an equilibrated Qiagen genomic tip 100 (Qiagen, Venlo, Netherlands). Genomic DNA was eluted and precipitated using isopropanol. Finally, HMW DNA was isolated by spooling.

### SMRTbell Library Preparation and Sequencing

HMW genomic DNA was sheared to 30 kb using a Megaruptor^®^ 2 instrument (Diagenode SA, Seraing, Belgium). DNA-damage and end repair was carried out prior to blunt adaptor ligation and exonuclease purification using ExoIII and ExoVII, in accordance with the protocol supplied by Pacific Biosciences (P/N 101-024-600-02, Pacific Biosciences, Menlo Park, CA). The resultant SMRTbell templates were size-selected using a BluePippin™ (Sage-Science, MA, USA) instrument with a 15 kb cut off and a 0.75% DF Marker S1 high-pass 15 kb −20 kb gel cassette. The final library was sequenced on a Sequel System (Pacific Biosciences) with v2 chemistry, MagBead loading and SMRT Link UI v4 analysis.

### Lucigen PCR-free Library Preparation and Sequencing

Genomic DNA was fragmented to 350 bp size using a Covaris S2 Focused Ultrasonicator (Covaris, Woburn, MA). Subsequent end-repair, A-tailing, Lucigen adaptor ligation, and size-selection was performed using the Lucigen NxSeq^®^ AMPFree Low DNA Library Kit (Lucigen, Middleton, WI). Libraries were quantified using a Qubit 2.0 instrument (Life Technologies, Carlsbad, CA) and library profiles were analyzed using a Bioanalyzer High Sensitivity Chip on an Agilent Bioanalyzer 2100 (Agilent Technologies, Santa Clara, CA). The libraries were sequenced to a coverage depth of 10X on an HiSeq 3000 instrument (Illumina) using a HiSeq 3000/4000 SBS kit and paired-end 150 base read chemistry.

### Genome assembly and haplotype merging

The genome was assembled from 58 Gb of Sequel long read data using *Canu* (version 1.6; genomeSize=544m; other parameters default) (41). Raw contigs were polished with *Arrow* (ConsensusCore2 version 3.0.0; consensus models S/P2-C2 and S/P2-C2/5.0; other parameters default) and *Pilon* (version 1.22; parameters default) (42). Polished contigs were repeat masked using *WindowMasker* (version 1.0.0; -checkdup; other parameters default) (43). Repeat-masked contigs were screened for misjoints and subjected to haplotype merging using *HaploMerger2* (commit 95f8589; identity=80, other parameters default (44). A custom scoring matrix was supplied to both *lastz* steps of *Haplomerger2* (misjoint and haplotype detection). The scoring matrix was inferred from an all-vs-all contig alignment using *minimap2* (version 2.10; preset asm10; other parameters default) (45) taking only the best contig-to-contig alignments into account. The final assembly was finished against the chromosome-resolved *A. hypochondriacus* genome (26) using *reveal finish* (commit 98d3ad1; –fixedgapsize –gapsize 15,000; other parameters default) (46). The 16 resulting pseudo chromosomes represented 99.6% of the original assembly.

### Alignment, SNP calling, and gene annotation

We used *freebayes* (47) to call SNPs jointly on all samples. For whole genome analyses, we used a thoroughly filtered SNP set following established guidelines (48, 49) adapted for whole genome data: sites were removed based on missing data (>80%), complexity, indels, allelic bias (<0.25 & >0.75), whether there was a discrepancy in paired status of reads supporting reference or alternate alleles, and mapping quality (QUAL < 30, representing sites with greater than a 1/1000 error rate). Individuals with excess missing data (>5%) were dropped. This led to a final, high confidence SNP set of 10,280,132 sites. For *EPSPS*-specific analyses and genome-wide investigations that required invariant sites, we recalled SNPs with *samtools* (V1.7; (50)) and *bwa-mem* (V0.7.17; (51)). For this snp set, sites were minimally filtered on mapping quality and missing data (keeping only sites with MQ > 30 & < 20% missing data), so that diversity estimates were not biased by preferentially retaining invariant or variant sites. For both snp sets, we used *bwa-mem* to map to our fastqs to the reference genome. Bam files were sorted and duplicates marked with *sambamba* (V0.6.6; (52)), while cigars were split and read groups added with *picard* (V2.17.11).

We performed gene annotation on both our final assembly and the *A. hypochondriacus*-finished pseudoassembly using the *MAKER* pipeline (53). *A. tuberculatus*-specific repeats were identified using *RepeatModeler* (v1.0.11; (13)), combined with the RepBase repeat library, and masked with *RepeatMasker* (v4.0.7; (14)). This repeat-masked genome was then run through *MAKER* (v2.31.8), using EST evidence from an *A. tuberculatus* transcriptome assembly (54) and protein homology evidence from *A. hypochondriacus* (55). The gene models were further annotated using *InterProScan* (v69.0; (56)), resulting in a total of 30,771 genes and 40,766 transcripts with a mean transcript length of 1,245 bp. The mean annotation edit distance (AED) score was 0.21, and 98.1% of the gene predictions had an AED score of <0.5, indicating high quality annotations.

### Phenotyping

Seedlings from each population were grown in a 1:1:1:1 soil:peat:Torpedo Sand:LC1 (SunGro commercial potting mix) medium supplemented with 13-13-13 Osmocote in a greenhouse that was maintained at 28°/22°C day/night temperatures for a 16:8 h photoperiod. Plants were sprayed at the 5-7 leaf stage with 1,260 g glyphosate (WeatherMax 4.5 L, Monsanto, Chesterfield, MO) per hectare. Fourteen days after treatment, plants were rated visually on a scale of 0 (highly sensitive) to 5 (no injury). Plants rated 2 or higher were classified as resistant. Prior to herbicide treatment, single leaf samples were taken from each plant and stored at −80°C until ready for gDNA extraction. Tissue from plants rated as highly glyphosate-resistant or susceptible were selected from each population for genomic DNA extraction using a modified CTAB method (40).

### Copy number estimates

Scaled coverage and copy number at *EPSPS* was estimated by dividing the coverage at each site across the focal region by the mode of genomewide coverage after excluding centromeric regions (which have repeats and thus often abnormally high coverage) and regions oflow coverage (<3X, indicative of technical coverage bias), which should represent the coverage of single-copy genes.

### Structure, demographic modelling & summary statistics

To model neutral demographic history and estimate neutral diversity, we used a Python script (available at https://github.com/tvkent/Degeneracy) to score 0-fold and 4-fold degenerate sites across the genome. This procedure estimated 17,454,116 0-fold and 4,316,850 4-fold sites across the genome, and after intersecting with our final high quality *freebayes*-called SNP set, resulted in 345,543 0-fold SNPs and 326,459 4-fold SNPs. The latter was used as input for demographic modelling.

Our two-population demographic model of *A. tuberculatus* modelled the split between the *A. tuberculatus* var. *tuberculatus* and var. *rudis* subspecies by collapsing individuals into one of the two populations based on predominant ancestry as identified in our *STRUCTURE* analyses, estimated in *∂*a*∂*i (V1.7.0) (57) using the pipeline available on https://github.com/dportik/dadi_pipeline (58). 1D and 2D site frequency spectrums were estimated using the program *easySFS* (https://github.com/isaacovercast/easySFS), and samples were projected downwards to maximize the number ofloci without missing data vs. number ofindividuals retained. We ensured that the log-likelihood of our parameter set had optimized by iterating the analysis over four rounds of increasing reps, from 10 to 40. We tested a set of 20 diversification models, with variation in split times, symmetry of migration, constancy of migration, population sizes and size changes. The most likely inferred demography followed a model of secondary contact, where initially populations split without gene flow, followed by population size change with asymmetrical gene flow, and included 8 parameters: Size of population 1 after split (nu1a), size of population 2 after split (nu2a), the scaled time between the split and the secondary contact (in units of 2*N_a_ generations) (T1). the scaled time between the secondary contact and present (T_2_), size of population 1 after time interval (nu1b), size of population 2 after time interval (nu2b), migration from population 2 to population 1 (2*N_a_*m12), and migration from population 1 to population 2 (m21). N_e_ was calculated by substituting the per-siteθestimate (after controlling for the effective sequence length to account for losses in the alignment and missed or filtered calls) and the *A. thaliana* mutation rate (7*10^−9^) (59) into the equation *θ* = 4*Neμ*.

We used *PLINK (V1.9; (60))* to perform a PCA of genotypes from our final *freebayes* SNP set after thinning to reduce the effects of sites that are in linkage disequilibrium, used STRUCTURE *(V2.3.4) (61)* to estimate admixture across populations, and *treemix (V3) (62)* to infer patterns of population splitting and migration events. To calculate summary statistics (*π*,F_ST_, *D_xy_*), we used scripts from the genomics general pipeline available at https://github.com/simonhmartin/genomics_general, binning SNPs into 100 kb windows with a step size of 10 kb. To estimate the proportion of introgression of var. *rudis* ancestry into Walpole agricultural populations in these genomic windows, we used the *f* statistic (but with non-overlapping windows, (28). For investigation ofintrogression of Natural populations into Illinois (var. *tuberculatus* into var. *rudis*), we used Missouri as the reference ingroup. For investigation ofintrogression of Essex populations into Walpole (var. *tuberculatus* into var. *rudis*), we used Natural populations as the reference in group. Lastly, for investigation of introgression of Natural populations into Essex (var. *tuberculatus* into var. *rudis*), we used Missouri as the reference ingroup. To get confidence intervals for the *f* statistic estimates, we performed jackknifing by calculating pseudovalues by removing one 250 kb block at a time.

For the outlier analysis of putative genes underlying contemporary agricultural adaptation in Walpole, we analyzed genome-wide differentiation (F_st_) in 10 kb windows, and classified windows as outliers when they were in the top 1% for extreme differentiation. A GO enrichment test was then performed for these outlier regions, after finding their intersecting annotated *A. tuberculatus* genes, and their orthologues in *A. thaliana* using *orthofinder (63)*. To look at the possibility of gene expansion in these enriched gene families, we first characterized normalized copy number in 50 SNP windows within each annotated gene in that family, for every individual. We then characterized the max copy number window within each gene for each individual, as heterogeneous mapping of paralogs/orthologs due to differential levels of degeneration should lead to variation in copy number across windows within the gene. We then took the average across all individuals for Walpole and Natural populations, and compared their distribution across all genes for each family. We tested whether the average of the max copy number window with in each gene differed consistently across all genes and between Walpole and Natural populations by performing an anova of region and gene ID, allowing for an interaction.

### Detecting selective sweeps and estimating recombination rate

To detect differences in the strength and breadth of sweep signal associated with selection from glyphosate across geographic regions, we used SNPs called from the pseudoassembly of our *A. tuberculatus* reference. Sweep detection can be strongly influenced by heterogeneity in recombination rate, and so as a control (in our *Sweepfinder2* and *XPEHH* analyses), we used the interval function in *LDhat* (64) to estimate variable recombination rate independently across all 16 chromosomes of the pseudoassembly, using a precomputed lookup table for aθof 0.01 for 192 chromosomes. Accordingly, we randomly subsetted individuals to retain only 96 individuals for computation of recombination rate estimates, which was implemented by segmenting the genome into 2,000 SNP windows, following the workflow outline in https://github.com/QuentinRougemont/LDhat_workflow.

To account for the fact that high-copy-number loci may allow for increased diversity relative to single copy regions, we randomly sampled one allele per locus along the length of chromosome 5 to create pseudo-haploid haplotypes for our sweep scans. This ensures that any increased differentiation is due to differences among individuals, rather than among haplotypes within individuals. The *XP-EHH* scan ((34), calculated based on the difference in haplotype homozygosity between amplified and nonamplified individuals for each geographic region after controlling for recombination rate, was implemented in *selscan* (65). Scripts available at https://github.com/simonhmartin/genomics_general were used for calculating differentiation and the difference in diversity. Pseudo-haploid haplotypes were also used to calculate a maximum likelihood tree for the 235 SNPs in *EPSPS*. For each tree, we realigned sequences before bootstrapping 1,000 replicates of our haplotree with *clustalomega* (66).In contrast to haplotype-based methods that required phased data, we also ran *Sweepfinder2* (32, 33), a program that compares the likelihood of a selective skew in the site frequency spectrum (SFS) at focal windows compared to the background SFS while controlling for heterogeneity in recombination rate. The SFSs of 10 kb windows across chromosome 5 were compared to the genome-wide SFSs at 4-fold degenerate sites, that for this analysis, was also randomly sampled for one allele per locus, for an equivalent comparison of the SFS. Lastly, we investigated similarity in the EPSPS amplification within and among populations and regions by estimating the Spearman’s rank correlation coefficient for all pairwise comparisons of resistant, amplification-containing individuals. This was done for the 1 Mb region surrounding EPSPS.

## Acknowledgments

We thank Tyler Kent & Anna O’Brien for useful discussion, and Rebecca Schwab, Fernando Rabanal and Talia Karasov for comments. We would also like to thank and recognize the Ojibwe, Potawatomi, and Odawa peoples of the Walpole Island First Nation. This work was supported by NSERC Discovery Grants (SIW, JRS), an NSERC PGS-D (JK), the Department of Ecology & Evolutionary Biology at University of Toronto (JK), IMPRS Molecules to Organisms (BW), Max Planck Society and Ministry for Science, and Research and Art of Baden-Württemberg in the Regio-Research-Alliance “Yield stability in dynamic environments” (DW). Author contributions Conceptualization: Julia M. Kreiner, Stephen I. Wright, John R. Stinchcombe, Detlef Weigel, Patrick J. Tranel Data curation: Felix Bemm, Bridgit Waithaka, Christa Lanz, Julia Hildebrandt, Julian Regalado, Darci Ann Giacomini Analysis: Julia M. Kreiner, Darci Ann Giacomini, Felix Bemm, Julian Regalado Funding acquisition: Detlef Weigel, Stephen I. Wright, John R. Stinchcombe, Patrick J. Tranel, Peter H.Sikkema, Julia M. Kreiner. Methodology: Julia M. Kreiner, Stephen I. Wright Supervision: Stephen I. Wright, John R. Stinchcombe, Patrick J. Tranel, Detlef Weigel Visualization: Julia M. Kreiner Writing – original draft: Julia M. Kreiner Writing – review & editing: Julia M. Kreiner, Stephen. I. Wright, John R. Stinchcombe, Detlef Weigel, Patrick J. Tranel, Peter H. Sikkema, Bridgit Waithaka

