## Supplementary material for "Multiple modes of convergent adaptation in the spread of glyphosate-resistant *Amaranthus tuberculatus*": Sup Figs and Tables

### Supplementary Tables and Figures

**Table S1. Metrics of raw and the haplotype-reduced reference genome assemblies.**

| Assembly | Total Size (Mb) | Sequence (#) | N50 (bp) | Longest sequence (bp) | Completeness (%) | Duplicates (%) |
|---|---|---|---|---|---|---|
| Raw Assembly | 1,159,758,700 | 4,207 | 905,938 | 9,167,955 | 90 | 71 |
| Haploid Assembly | 663,660,067 | 2,514 | 1,738,871 | 13,655,724 | 87 | 6 |

**Table S2. Correlation of PC1 and PC2 (Fig. 1E) with both longitude and latitude.** From 4 separately run ANOVAs.

| | | | |
|---|---|---|---|
| **Longitude** | | | |
| **PC1** | $r^2 = 0.05279$ | $p = 0.00190$ | $F_{1,160} = 9.973$ |
| **PC2** | $r^2 = 0.7685$ | $p = <2.2e^{-16}$ | $F_{1,160} = 535.3$ |
| **Latitude** | | | |
| **PC1** | $r^2 = 0.08058$ | $p = 0.0001482$ | $F_{1,160} = 15.11$ |
| **PC2** | $r^2 = 0.6025$ | $p = <2.2e^{-16}$ | $F_{1,160} = 245.1$ |

**Table S3.** Copy number of genes in the cytochrome p450 and glycosyltransferase families that significantly positively relate to phenotypic resistance ratings, after controlling for multiple testing.

| **Glycosyltransferase** | ***F*** | **raw *p* value** | **holm-corr. *p*** |
|---|---|---|---|
| Adenine DNA glycosylase | 16.2 | 0.0002 | 0.01 |
| UDP-glycosyltransferase 87A2 | 13.4 | 0.0007 | 0.04 |
| xyloglucan glycosyltransferase 6 | 14.2 | 0.0005 | 0.03 |
| **CYP450** | | | |
| Cytochrome P450 734A1 | 15.6 | 0.0003 | 0.02 |
| Cytochrome P450 71D9 | 22.5 | 0.00002 | 0.002 |

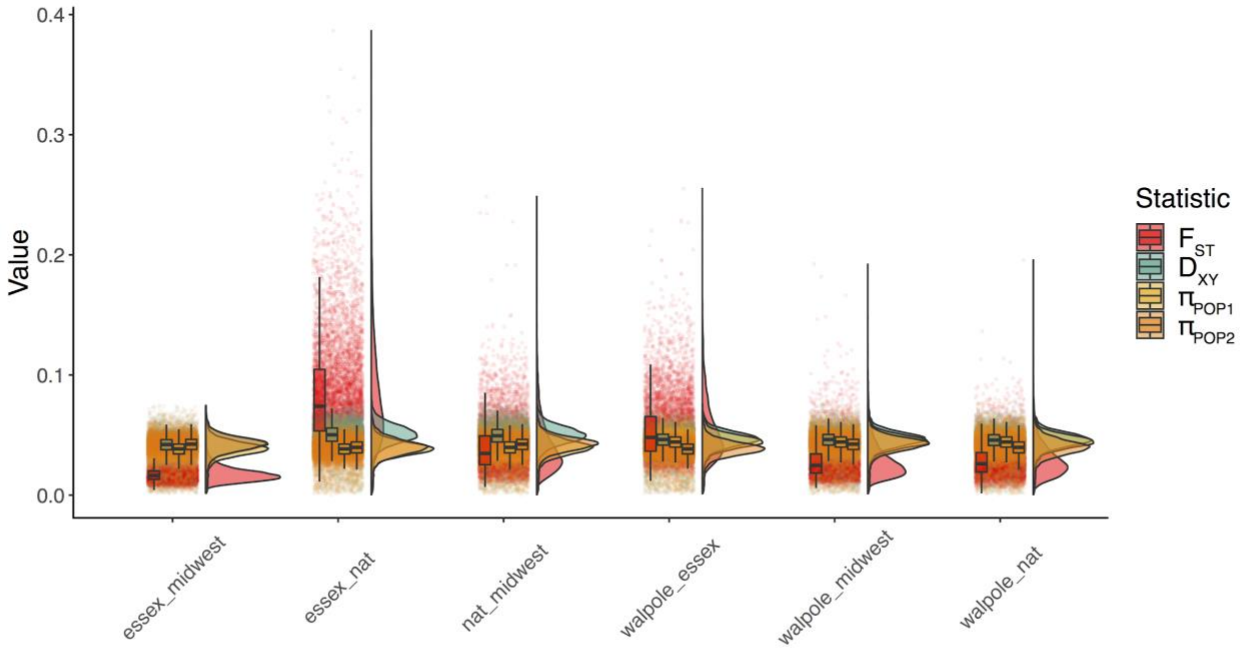


**Fig. S1.** The relationship between absolute diversity ($D_{xy}$), relative diversity ($F_{st}$), and within-population diversities ($\pi$) for among geographic region comparisons. For each comparison, points refer to the means of 100 kb windows, with corresponding boxplots and density curves for each summary statistic. Pop1 in the legend refers to the first population of the pairwise comparison, and Pop2 to the second.

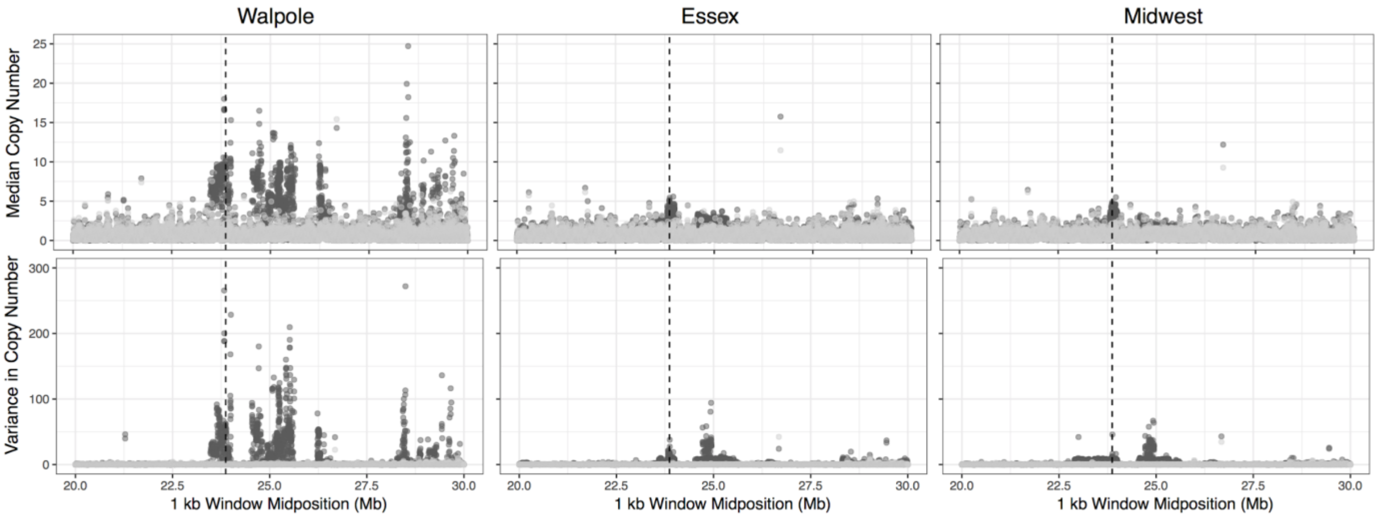


**Fig. S2.** Median and variance in *EPSPS* copy number for individuals with (dark grey) and without (light grey) copy number increase in chromosome 5. Dashed vertical lines indicate location of *EPSPS*.

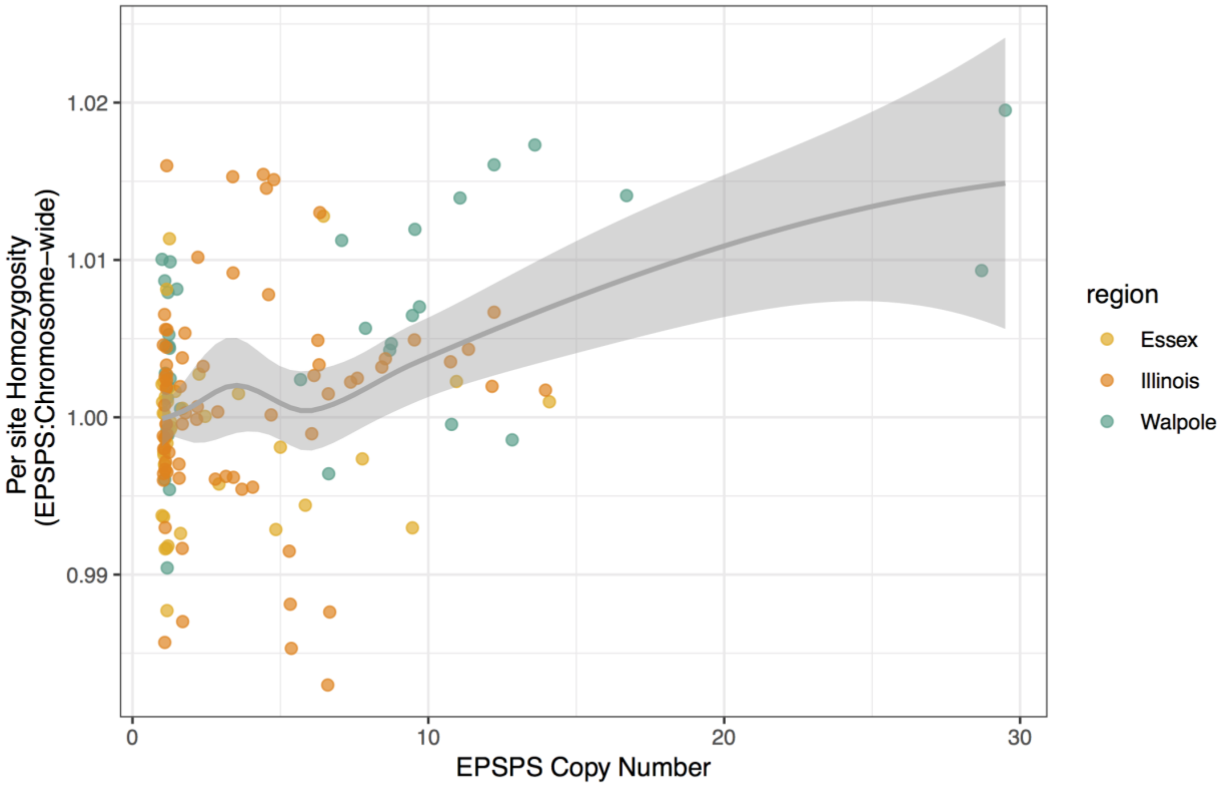


**Fig. S3.** The relationship between the ratio of *EPSPS* copy number and per-site heterozygosity in 1 Mb around EPSPS relative to all of chromosome 5. An ANOVA for ratio of heterozygosity to EPSPS copy number was significant ($p$ = 5.2e-07 and $r^2$ = 0.147).

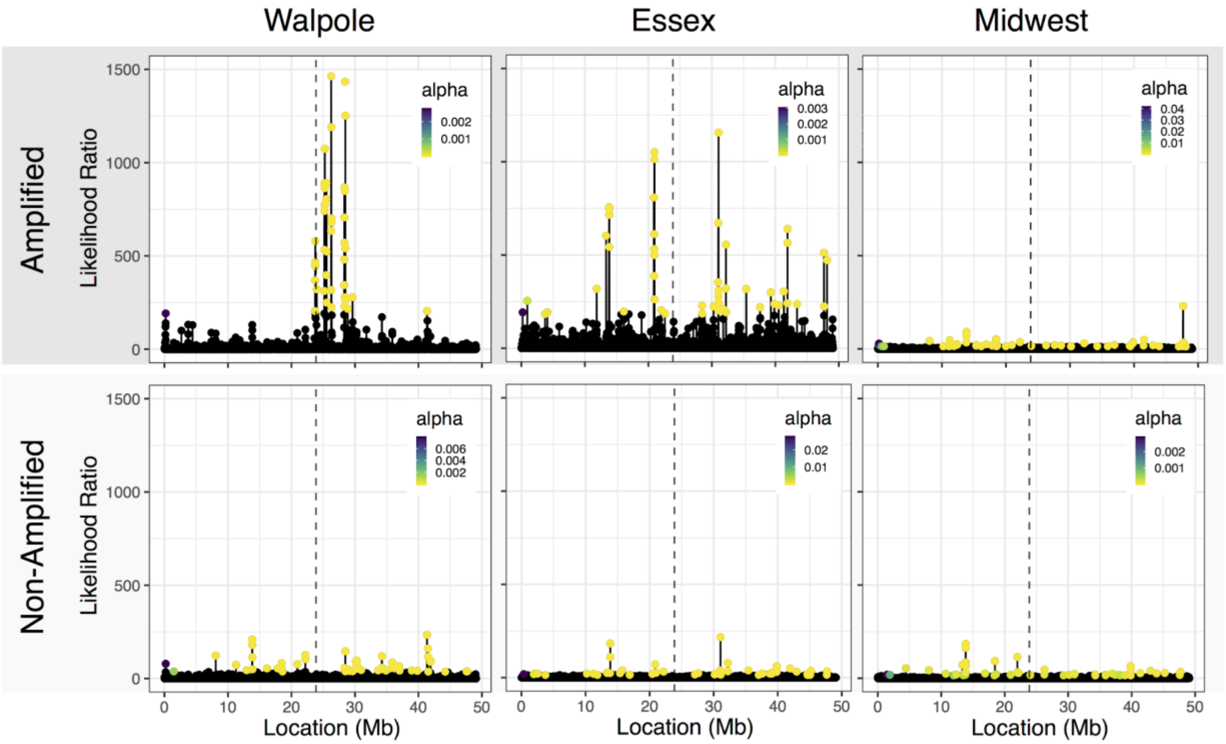


**Fig. S4.** *Sweepfinder2* likelihood ratio scores of a sweep occurring across chromosome 5. Scores were controlled for recombination rate variation and the genome-wide neutral site frequency spectrum. Alpha refers to the relative strength of recombination vs. selection, with smaller values indicating stronger selection (31).

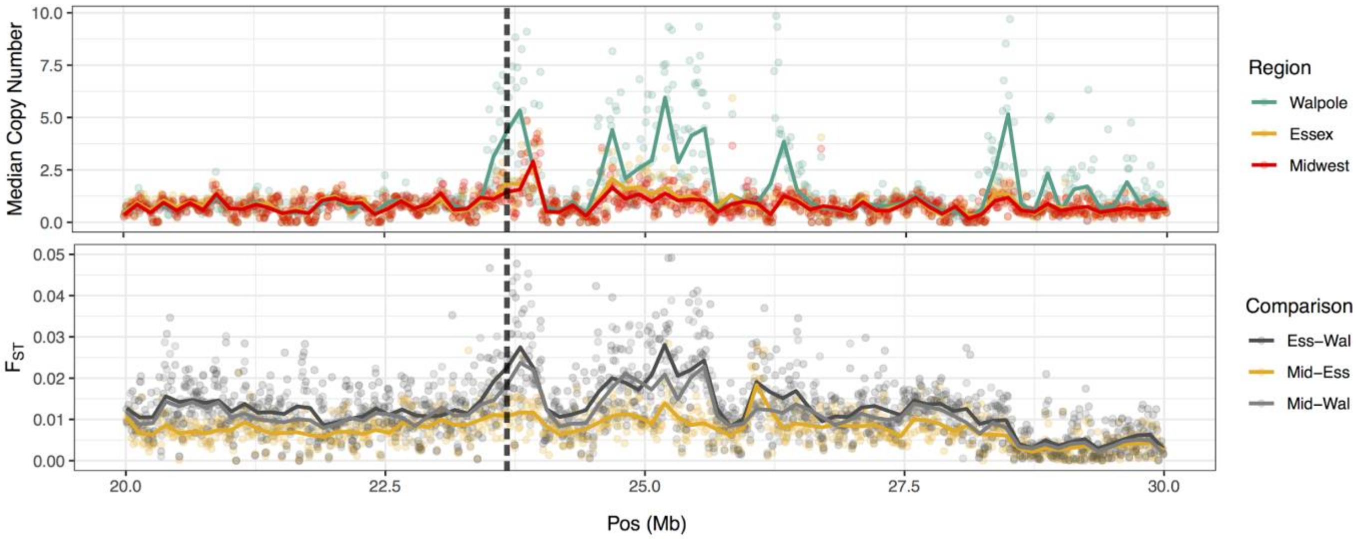


**Fig. S5.** Genetic differentiation ($F_{ST}$) between agricultural regions and median copy number within agricultural regions in 10kb windows along the 20 Mb - 30 Mb region of chromosome 5. *EPSPS* indicated by the vertical dashed grey black line.
